# Cryo-EM Structures of Amyloid-β Filaments With the Arctic Mutation (E22G) From Human and Mouse Brains

**DOI:** 10.1101/2022.11.07.515410

**Authors:** Yang Yang, Wenjuan Zhang, Alexey G. Murzin, Manuel Schweighauser, Melissa Huang, Sofia Lövestam, Sew Y. Peak-Chew, Takashi Saito, Takaomi C. Saido, Jennifer Macdonald, Isabelle Lavenir, Bernardino Ghetti, Caroline Graff, Amit Kumar, Agneta Nordberg, Michel Goedert, Sjors H.W. Scheres

## Abstract

The Arctic mutation, encoding E693G in the *amyloid precursor protein (APP)* gene [E22G in amyloid-β (Aβ)], causes dominantly inherited Alzheimer’s disease. Here we report the high-resolution cryo-EM structures of Aβ filaments from the frontal cortex of a previously described case (*AβPParc1*) with the Arctic mutation. Most filaments consist of two pairs of non-identical protofilaments that comprise residues V12-V40 (human Arctic fold A) and E11-G37 (human Arctic fold B). They have a substructure (residues F20-G37) in common with the folds of type I and type II Aβ42. When compared to the structures of wild-type Aβ42 filaments, there are subtle conformational changes in the human Arctic folds, because of the lack of a side chain at G22, which may strengthen hydrogen bonding between mutant Aβ molecules and promote filament formation. A minority of Aβ42 filaments of type II was also present, as were tau paired helical filaments. In addition, we report the cryo-EM structures of Aβ filaments with the Arctic mutation from mouse knock-in line *App*^*NL-G-F*^. Most filaments are made of two identical mutant protofilaments that extend from D1-G37 (murine Arctic fold). In a minority of filaments, two dimeric folds pack against each other in an anti-parallel fashion. The murine Arctic fold differs from the human Arctic folds, but shares some substructure.

## INTRODUCTION

Dominantly inherited mutations in the amyloid precursor protein gene (*APP*) that cause disease are a mainstay of the amyloid cascade hypothesis of Alzheimer’s disease (AD) (11,14). They have shown that overexpression of wild-type amyloid-β peptide (Aβ) or an increase in the ratio of Aβ peptides of 42 to 40 residues (Aβ42/Aβ40) is sufficient to cause familial AD. We recently reported that Aβ42 filaments from sporadic and inherited (*APP*^V717F^ and *PSEN1*^F105L^) cases of AD share identical structures (35). These mutations lead to the deposition of wild-type Aβ42.

Several mutations in *APP* give rise to mutant Aβ, including dominantly inherited *APP*^E693G^ (Arctic mutation) (17,24), *APP*^E693K^ (Italian mutation) (4) and *APP*^E693Q^ (Dutch mutation) (21), as well as recessively inherited *APP*^ΔE693^ (Osaka mutation) (33). The structures of mutant Aβ filaments from brain are not known. Mutations E693K and E693Q give rise to cerebral amyloid angiopathy, resulting in focal symptoms related to recurrent strokes (3,5). Many mutation carriers also develop dementia, which often follows the strokes. Aβ deposits are more abundant in cerebral blood vessels than in brain parenchyma and there are no abundant neuritic plaques or tau inclusions.

In contrast, mutations E693G and ΔE693 cause early-onset AD. For the Arctic mutation, unlike in sporadic AD, symptomatic carriers are negative for Pittsburgh compound B (PiB) by positron emission tomography (PET) (28). Like in sporadic AD, they show reduced Aβ42 and elevated total tau and P-tau in cerebrospinal fluid, as well as cerebral hypometabolism, measured by ^18^F-fluorodeoxyglucose PET (20,25,32). By immunohistochemistry, amyloid plaques appear to be ring-shaped, contain truncated Aβ40 and Aβ42, and lack a congophilic core (1,16). A ring shape is observed predominantly with Aβ42 antibodies, with Aβ40 staining being distributed more homogenously through plaques (20).

The Arctic mutation causes diminished rather than increased levels of Aβ40 and Aβ42 in conditioned media from transfected cells (30). This paradox has been explained by the finding that E22G Aβ40 forms what has been described as protofibrils at a faster rate and in larger number than wild-type Aβ40 (24). Increased assembly of Aβ with the Arctic mutation compared to wild-type was greater for Aβ40 than for Aβ42 (22). Based on results with the Arctic mutation and other findings, it has been proposed that the neurotoxic effects of Aβ are mainly mediated by oligomers and protofibrils rather than filaments (4,19,34). In cases of AD with the Arctic mutation, tau pathology was mainly in the form of neuropil threads, but neurofibrillary tangles and neuritic plaques were also present (16).

Here we show that the structures of mutant Aβ filaments from the frontal cortex of an individual with missense mutation E693G in *APP* differ from those of wild-type Aβ filaments. We also show that the structures of Aβ filaments from mice knock-in for the Arctic mutation (line *App*^NL-G-F^) (26) differ from those present in human brains, raising doubts about the relevance of this mouse model for studying AD.

## MATERIALS AND METHODS

### Genetics and clinical history

We determined the cryo-EM structures of Aβ and tau filaments from the frontal cortex of a previously described female individual (*AβPParc1*) with mutation E693G in APP (20,28). The presence of a heterozygous *APP* Arctic mutation (c.2078A>G) in exon 17 was confirmed by re-sequencing of DNA extracted from the blood of the tissue donor. AmpliTaqGold 360 PCR Master Mix (Thermo Fisher Scientific) was used for PCR amplification. Primer sequences and PCR conditions are available upon request. Sanger sequencing in both directions was performed using the BigDye Terminator v3.1 cycle sequencing kit (Thermo Fisher Scientific) and analyzed using an ABI3500 genetic analyzer. As reported (28), the proband began to experience cognitive symptoms at the age of 53 years, was diagnosed with AD at age 62 and died aged 66. The neuropathological characteristics of case *AβPParc1* have been described (28).

### *App*^*NL-G-F*^ knock-in mice

We determined the cryo-EM structures of Aβ filaments from the brain of a 22-month-old homozygous *App*^*NL-G-F*^ knock-in mouse on a C57 BL/6 JAX background. Beginning at 2 months of age, these mice form abundant extracellular deposits that are made of human Aβ with Arctic mutation E22G (26). They carry the Swedish double mutation (KM670/671NL), the Arctic mutation (E693G) [E22G in humanized Aβ] and the Beyreuther/Iberian mutation (I716F) in APP.

### Extraction of filaments

For cryo-EM analysis of the human sample, sarkosyl-insoluble material was extracted from temporal cortex of case *AβAPParc1*, essentially as described (31). Briefly, tissues were homogenized in 20 vol (w/v) extraction buffer consisting of 10 mM Tris-HCl, pH 7.4, 0.8 M NaCl, 10-20% sucrose and 1 mM EGTA. Homogenates were brought to 2% sarkosyl and incubated for 60 min at 37° C. Following a 10 min centrifugation at 10,000 g, the supernatants were spun at 100,000 g for 60 min. The final pellets were resuspended in 100 µl/g of 20 mM Tris-HCl, pH 7.4, 50 mM NaCl. For cryo-EM analysis of the mouse samples, sarkosyl-insoluble material was extracted from whole brains of mouse knock-in line *App*^*NL-G-F*^. Tissues were homogenized in 20 vol (w/v) extraction buffer consisting of 20 mM Tris-HCl, pH 7.4, 0.8 M NaCl, 15% sucrose, 5 mM EGTA, 1% sarkosyl and protease inhibitor (1 tablet per 10 ml, Roche), and incubated for 60 min at room temperature. Following a 20 min centrifugation at 10,000 g, the supernatants were spun at 124,000 g for 45 min at 20° C. The pellets were resuspended in 200 µl extraction buffer, followed by a second spin at 124,000 g. Pellets were then resuspended as above, followed by a third spin at 124,000 g. The final pellets were resuspended in 33-100 µl/g of 20 mM Tris-HCl, pH 7.4, 200 mM NaCl and used for cryo-EM analysis.

### Mass spectrometry

Mass spectrometry was performed as described (23). Sarkosyl-insoluble pellets were resuspended in 1ml/g extraction buffer and centrifuged at 3,000 g for 5 min. The supernatants were diluted 3-fold in 50 mM Tris-HCl, pH 7.4, containing 0.15 M NaCl, 10% sucrose and 0.2% sarkosyl, and spun at 100,000 g for 60 min. The pellets were resuspended in 100 µl hexafluoroisopropanol. Following a 3 min sonication at 50% amplitude (QSonica), they were incubated at 37° C for 2 h and centrifuged at 100,000 for 15 min, before being dried by vacuum centrifugation. Matrix-assisted laser desorption/ionization time of flight (MALDI-TOF) mass spectrometry was carried out as described (35).

### Electron cryo-microscopy

For cryo-EM, extracted Aβ filaments were centrifuged at 3,000 g for 2 min and treated with 0.4 mg/ml pronase for 30-60 min (12). Holey carbon grids (Quantifoil AuR1.2/1.3, 300 mesh) were glow-discharged with an Edwards (S150B) sputter coater at 30 mA for 30 s. Three µl aliquots were applied to the grids and blotted for 3-5 s with filter paper at 100% humidity and 4° C using a Thermo Fisher Vitrobot Mark IV. Datasets were acquired on Thermo Fisher G2 and G3 microscopes, with Gatan K3 detectors in counting mode, using a Bioquantum energy filter (Gatan) with a slit width of 20 e^−^V. Images were recorded with a total dose of 40 electrons per Å^2^.

### Helical reconstruction

All super-resolution frames were gain-corrected, binned by a factor of 2, aligned, dose-weighted and then summed into a single micrograph using RELION’s own implementation of MotionCor2 (38). Contrast transfer function (CTF) parameters were estimated using CTFFIND-4.1 (37). Subsequent image-processing steps were performed using helical reconstruction methods in RELION (15,39). Filaments were picked manually [dataset from frontal cortex of human Arctic case] or automatically using Topaz in RELION [dataset from brains of *App*^NL-G-F^ knock-in mice] (2). Reference-free 2D classification was performed to identify homogeneous segments for further processing. Initial 3D reference models were reconstructed *de novo* from 2D class averages (26) using an estimated rise of 4.75 Å and helical twists according to the observed cross-over distances of filaments in the micrographs. In order to increase the resolution of the reconstructions, Bayesian polishing and CTF refinement were performed (40). Final 3D reconstructions, after auto-refinement, were sharpened using the standard post-processing procedures in RELION, and resolutions calculated from Fourier shell correlations at 0.143 between the two independently refined half-maps, using phase-randomisation to correct for convolution effects of a generous, soft-edged solvent mask. Further details of data acquisition and processing are given in Table S1.

### Model building and refinement

Atomic models were built manually in Coot (7). Coordinate refinements were performed using *Servalcat* (36). Final models were obtained using refinement of only the asymmetric unit against the half-maps in *Servalcat*.

## RESULTS

### Structure of Aβ filaments from a human case with the Arctic mutation

We determined the cryo-EM structures of Aβ filaments from a case with mutation E693G in APP [E22G in Aβ] (Figure 1a). Most filaments, solved to 2.0 Å resolution, are made of four mutant protofilaments, with two copies each of non-identical protofilaments A and B (Figure 1b). The ordered cores of protofilaments A and B, hereafter referred to as the human Arctic folds A and B, consist of residues V12-V40 and E11-G37, respectively. Each fold comprises four β-strands (β1–β4) that extend between residues 12-15, 18–21, 30–32 and 34-36 in fold A, and between residues 11-13, 14-19, 30-32 and 34-36 in fold B (Figure 1c). Human Arctic fold A is almost identical to the fold of type II protofilaments of wild-type Aβ42, but is shorter by two C-terminal amino acids (Figure 1d). Human Arctic fold B is three amino acids shorter at its C-terminus than fold A, with residues F20-G37 adopting an almost identical conformation to that of fold A. Segment E11-F19 of fold B is one amino acid longer than that of fold A and adopts a different conformation.

**Figure 1.**
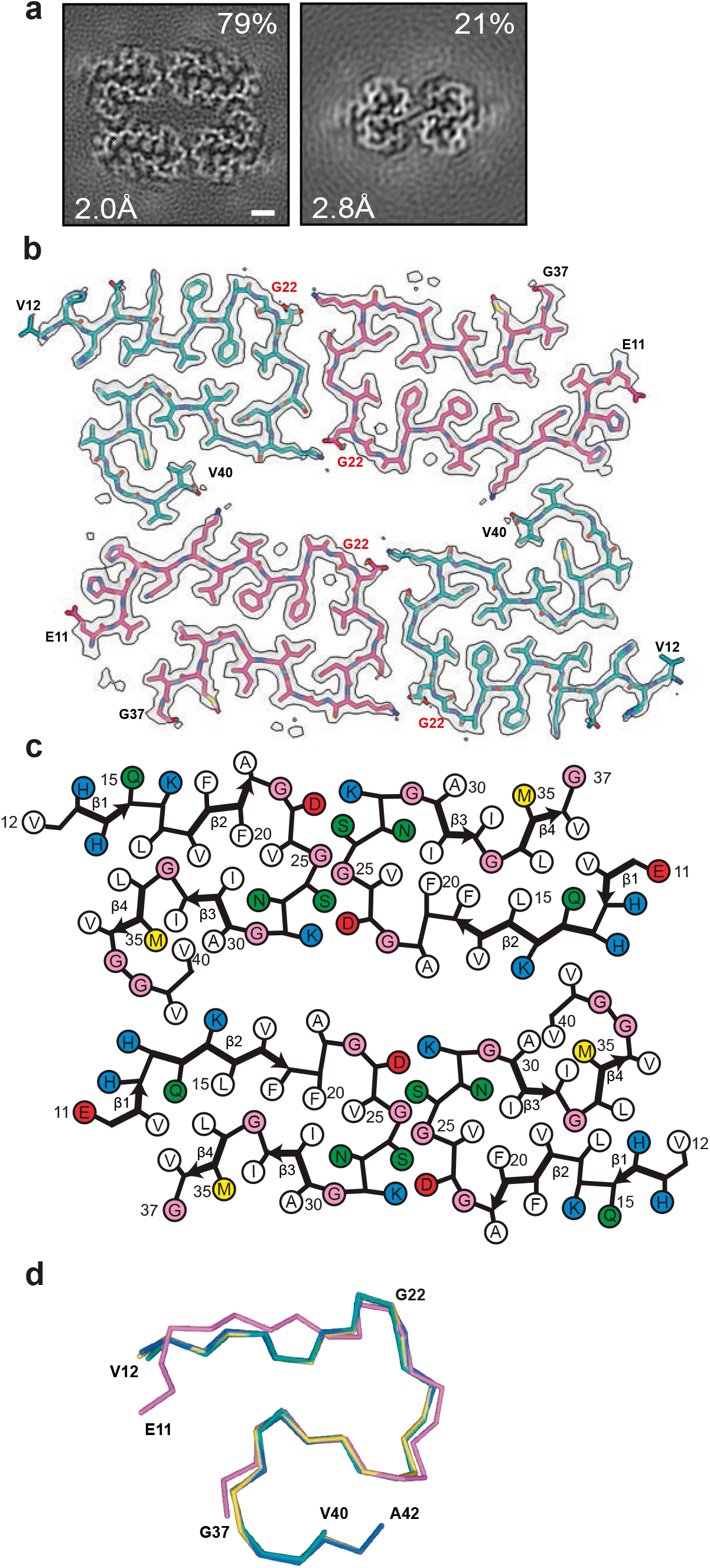
The human Arctic folds of E22 Aβ. a. Cross-sections of Aβ filaments from the frontal cortex of case *AβPParc1* perpendicular to the helical axis, with a projected thickness of approximately one rung. Percentages of filaments (relative to the total, taken as 100%) are shown on the top right. The resolutions of the cryo-EM maps are given on the bottom left (2.0 and 2.8 Å). Scale bar, 1 nm. b. Cryo-EM density maps (in transparent grey) and atomic models of the human Arctic folds. Human Arctic fold A (cyan) and human Arctic fold B (magenta). c. Schematic of human Aβ filaments with the Arctic mutation. Negatively charged residues are shown in red, positively charged residues in blue, polar residues in green, non-polar residues in white, sulphur-containing residues in yellow and glycines in pink. Thick connecting lines with arrowheads indicate β-strands. d. Superposition of the backbone structures of human Arctic fold A (cyan), human Arctic fold B (magenta), human Arctic type II Aβ42 protofilament (yellow) and human wild-type type II Aβ42 protofilament (blue). The all-atom r.m.s.d. values for human Arctic fold A with human Arctic fold B (residues F20-G37), human Arctic type II Aβ42 (residues V12-V40) and human wild-type type II Aβ42 structures (residues V12-V40) were 2.21 Å, 0.34 Å and 0.31 Å, respectively.

The substructures that are common to folds A and B form a dimeric and pseudo-symmetric interface that is centred on residues G25 and S26 from both folds and is stabilized by salt bridges between D23 from one fold and K28 from the other fold, and vice versa. These doublets of protofilaments A and B pack together with C2 symmetry to form the tetrameric filaments. The interface between doublets is also stabilized by two salt bridges, between the main chain carboxyl group of the C-terminal residue V40 from protofilament A in one doublet and the side chain of K16 from protofilament B in the other doublet, and vice versa (Figure 1b).

Besides tetrameric Aβ filaments, a minority of dimeric type II Aβ42 filaments was also present (Figure 1a). Solved to 2.8 Å resolution, their structures suggest that they are also made of mutant protein, but the presence of some wild-type Aβ42 cannot be excluded. Mass spectrometry of sarkosyl-insoluble fractions showed abundant mutant Aβ40 that was sequentially truncated at the N-terminus, and a smaller amount of wild-type Aβ42 (Figure S1a). There was also a substantial amount of mutant Aβ species truncated at E3 or E11, with these glutamate residues having been converted to pyroglutamates.

The Arctic mutation lies in the substructure that is shared by wild-type and mutant Aβ folds. Our high-resolution cryo-EM maps of Aβ filaments with the Arctic mutation showed the absence of side chain densities at G22, unlike the corresponding maps for wild-type Aβ filaments with side chain densities at E22 (Figure 2a-c). In filaments made of wild-type Aβ, the presence of a side chain at E22 restricts the orientation of the flanking main chain peptide groups and prevents the formation of hydrogen bonds linking these groups to those in other Aβ molecules. Removal of the side chains by the E22G mutation results in a slight reorientation of peptide groups in the “frustrated” loop F20-V24, which leads to increased hydrogen bonding between adjacent Aβ molecules (Figure 2d,e).

**Figure 2.**
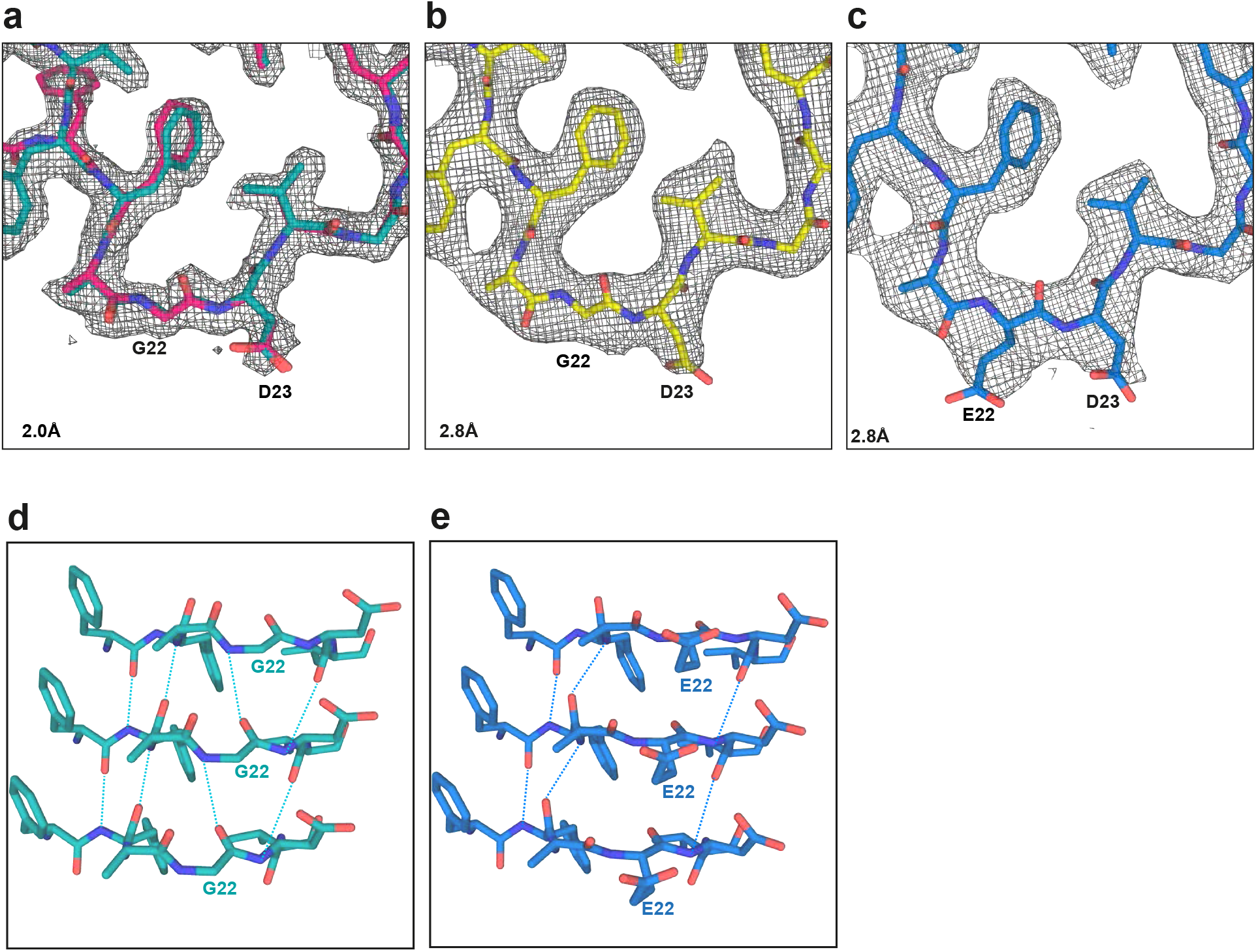
Structures of the E22G site in human Arctic and type II Aβ42 filaments. a. Structure of the F20-V24 arc of human Arctic fold A (cyan) superimposed on that of human Arctic fold B (magenta), and overlaid on the corresponding section of the 2.0 Å electron density map (grey). b. Structure of the F20-V24 arc of E22G type II Aβ42 fold (yellow) overlaid on the corresponding section of the 2.8 Å electron density map (grey). c. Structure of the F20-V24 arc of wild-type type II Aβ42 fold (blue) overlaid on the corresponding section of the 2.8 Å electron density map (grey). d. Side view of structure of human Arctic fold A G22 (cyan), showing the presence of hydrogen bonds (dashed lines) between the main chain groups. e. Side view of structure of human wild-type type II Aβ42 fold A E22 (blue), showing the presence of hydrogen bonds (dashed lines) between the main chain groups.

### Arctic fold of Aβ from *App*^*NL-G-F*^ mouse brains

*App*^*NL-G-F*^ knock-in mice deposit Aβ faster than *App*^*NL-F*^ knock-in mice (26). The cryo-EM structures of Aβ42 filaments from the brains of homozygous *App*^*NL-F*^ mice, which showed the type II Aβ42 fold, have been described (35). We now determined the cryo-EM structure of Aβ filaments from the brains of *App*^*NL-G-F*^ mice to 3.5 Å resolution (Figure 3). Filaments are made of two identical S-shaped mutant protofilaments that extend from D1-G37 of Aβ (Figure 3a,b). Each protofilament consists of five β-strands that extend from residues 1-8, 10–16, 17-19, 30–32 and 34-36 (Figure 3c). Additional densities around K16 may be co-factors or post-translational modifications of mutant Aβ, but their chemical identity remains unknown. We name the conformation of these protofilaments the murine Arctic fold.

**Figure 3.**
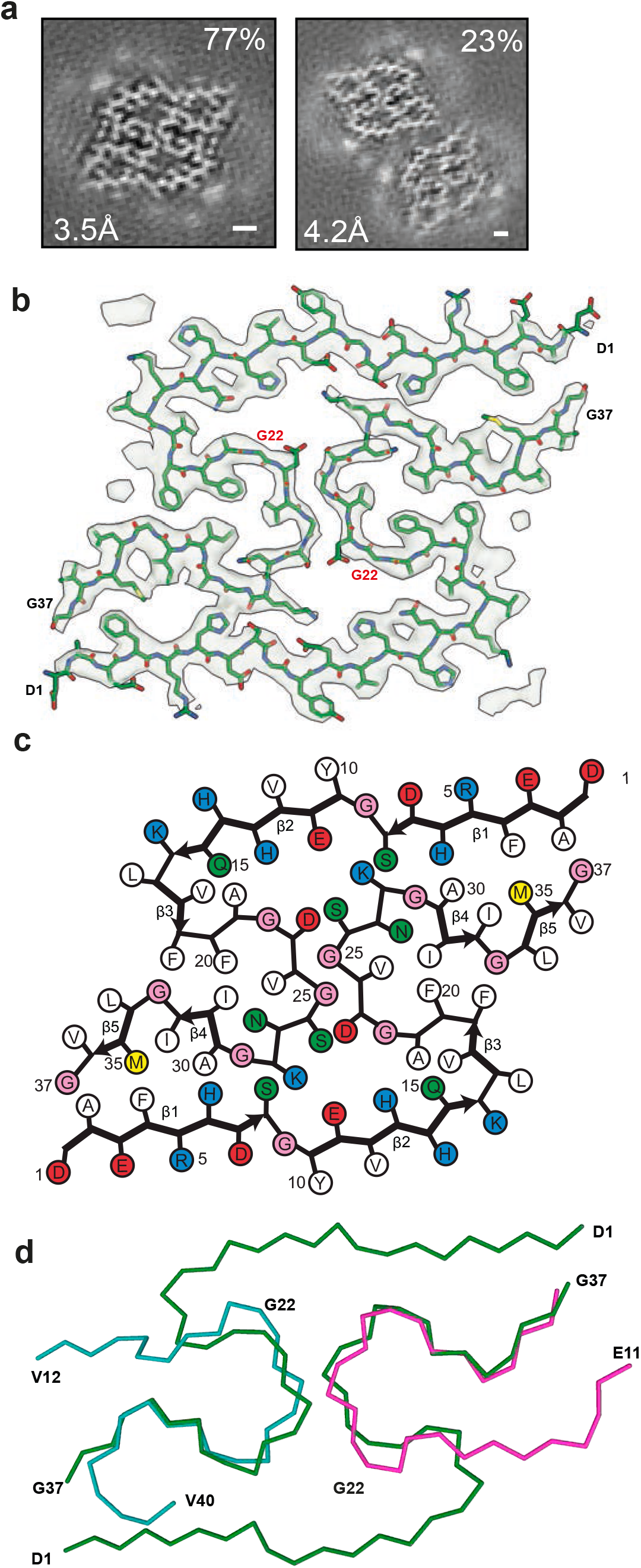
The murine Arctic fold of E22 Aβ. a. Cross-sections of Aβ filaments from the brains of *App*^NL-G-F^ mice perpendicular to the helical axis, with a projected thickness of approximately one rung. Percentages of filaments (relative to the total, taken as 100%) are shown on the top right. The resolutions of the cryo-EM maps are given on the bottom left (3.5 Å and 4.2 Å). Scale bar, 1 nm. b. Cryo-EM density map (in transparent grey) and atomic model (in green) of murine Aβ filaments with the Arctic fold. c. Schematic of murine Aβ filaments with the Arctic mutation. Negatively charged residues are shown in red, positively charged residues in blue, polar residues in green, non-polar residues in white, sulphur-containing residues in yellow and glycines in pink. Thick connecting lines with arrowheads indicate β-strands. d. Superposition of the backbone structures of dimeric murine Arctic filament (green) and the doublet of human Arctic fold A (cyan) and human Arctic fold B (magenta). The all-atom r.m.s.d. value for pairs of common substructures (F20-G37) of human Arctic folds A and B and murine dimeric Arctic fold was 2.35 Å.

It shares the substructure F20-G37 with human Arctic folds A and B, in which the backbone conformation of the loop F20-V24 differs from human Arctic folds in that the orientation of the peptide group between A21 and G22 is reversed. Flipping the peptide group does not affect its ability to form additional hydrogen bonds with other Aβ molecules, but places G22 in the glycine-only quadrant of the Ramachandran plot. The N-terminal segment (residues D1-F19) is longer than in human Arctic folds A and B, allowing it to fold back on the common substructure and extend across the dimeric interface towards the other protofilament (Figure 3d). Both protofilaments pack against each other with pseudo-2^1^ symmetry. The central portion of their interface is made of residues D23, G25 and S26 from both protofilaments and resembles the doublet-forming interface of human Arctic protofilaments A and B. At the edges, both protofilaments pack against each other through the hydrophobic side chains of A2 and F4 from one protofilament and A30, I31 and M35 from the other.

In addition, there are salt bridges between E11 and K28, and close contacts of the side chains of H6 and S8 with the backbone atoms of G29 and K28, respectively. We also observed a minority of wider filaments, in which two dimeric folds pack against each other in an anti-parallel fashion (Figure 3a). From the cryo-EM maps, we only observed residues ranging from D1 to G37 of mutant Aβ, consistent with mass spectrometry, which indicated that most Aβ in the sample is mutant Aβ1-38 (Figure S1b). It will be interesting to stain *App*^NL-G-F^ brains with antibodies specific for C-terminally truncated Aβ

### Tau filament structures from case *AβPParc1*

Tau filaments were found in the sarkosyl-insoluble fractions from frontal cortex of case *AβPParc1*. Their cryo-EM structures were determined to a resolution of 2.9 Å and found to be identical to those of PHFs from AD and some other diseases (Figure 4) (29). Straight tau filaments were not observed.

**Figure 4.**
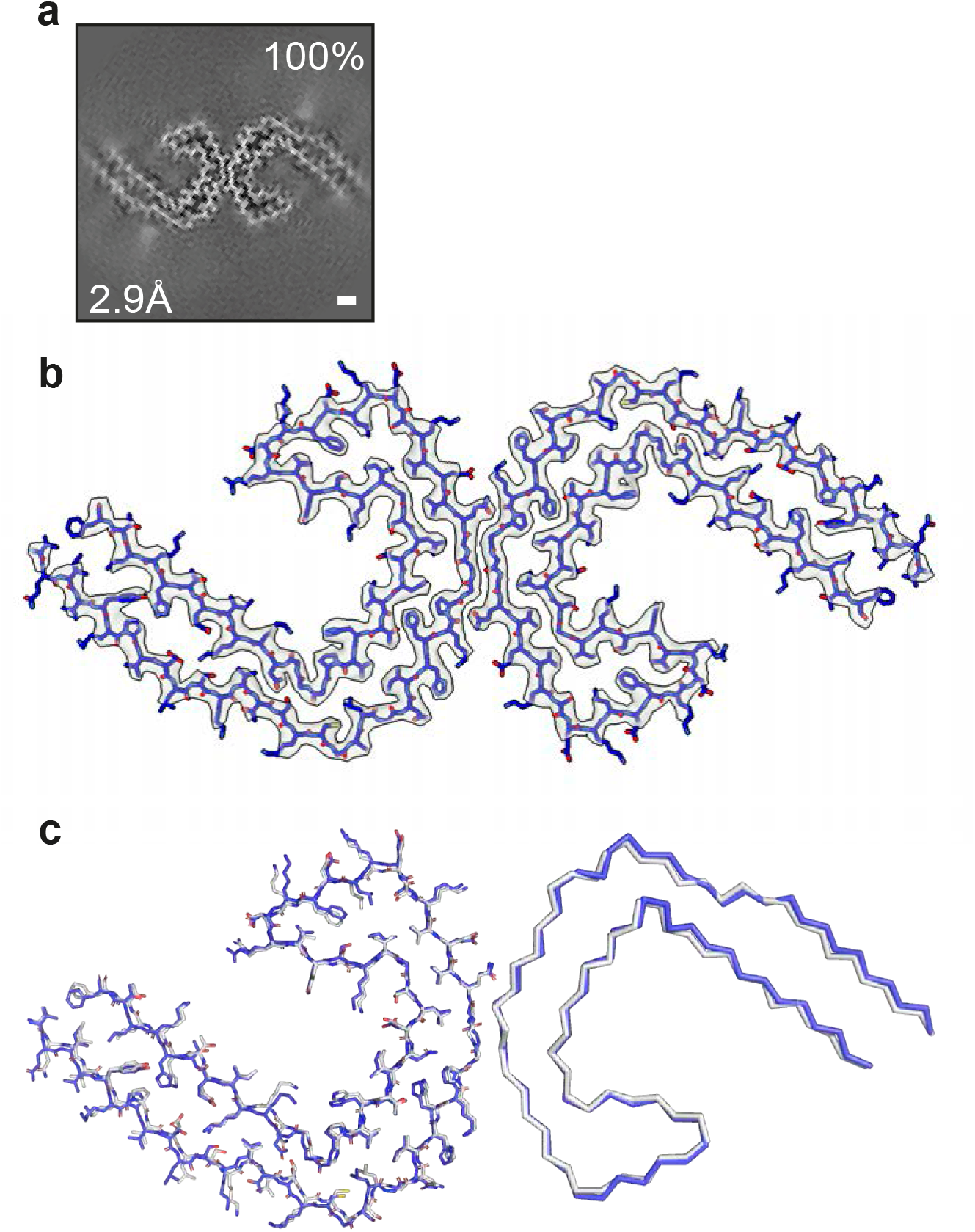
PHF tau filaments from case *AβPParc1*. a. Cross-sections of tau filaments from the frontal cortex perpendicular to the helical axis, with a projected thickness of approximately one rung. Percentage of filaments (relative to the total taken as 100%) are shown on the top right. The estimated resolution of the cryo-EM map is given on the bottom left (2.9 Å). b. Cryo-EM density map (grey) and atomic model of PHF (blue) from a human case with the Arctic mutation. c. Comparison of the PHF structure from case *AβPParc1* (blue) with the PHF structure from sporadic Alzheimer’s disease brain (grey) (PDB 5O3L). The structures are shown as sticks for one protofilament and as ribbons for the other protofilament.

## DISCUSSION

We report the first structures of filaments made of human mutant Aβ from brain. Tetrameric filaments containing the E22G Arctic mutation differ from dimeric type I and type II filaments of wild-type Aβ42. However, there is a large common substructure that is shared between protofilaments. Comparison of local conformations in this region revealed the presence of additional hydrogen bonds between adjacent Aβ molecules in mutant protofilaments; these hydrogen bonds cannot form in wild-type protofilaments, providing a plausible explanation for the increased fibrillogenesis of E22G Aβ. Mutation E22G may have an additional fibrillation-promoting effect, which is the relief of electrostatic repulsion as a result of removal of the negatively charged carboxylic group of E22; in the wild-type structure, E22 is trapped in close proximity to the carboxylic group of D23 (35).

These findings also suggest that the differences in structure between wild-type and mutant filaments may result from the E693G mutation affecting the cleavage of APP. Indeed, C2-symmetric mutant Aβ filaments comprise two asymmetric doublets, with protofilaments A and B adopting different folds. Human Arctic fold A ends strictly at V40, whereas Arctic fold B ends at G37, but may contain additional residues in its fuzzy coat. Mass spectrometry of sarkosyl-insoluble material indicates the presence of an N-terminal fuzzy coat of Aβ and shows that most mutant peptides end at residue V40. Similar mass spectrometry results have been reported from temporal cortex of another individual with the Arctic mutation (25). In agreement with these observations, immunohistochemistry of cerebral cortex from case *AβPParc1* has shown stronger staining for Aβ40 than Aβ42 (20). Previously, the structures of wild-type Aβ40 filaments were reported from the meninges of Alzheimer’s disease patients (18). The Arctic folds are different from these structures.

A minority of type II Aβ42 filaments was also observed. Apart from a lack of side chain density at G22, they are identical in structure to wild-type Aβ42 filaments (35). Co-deposition of wild-type Aβ and E22GAβ has been described (25), suggesting a possible link between their assemblies. Our results further suggest that they might co-assemble within the same filaments. Incorporation of a small amount of wild-type Aβ into E22G type II Aβ42 filaments would be invisible in the cryo-EM density maps.

The large common substructures shared by human Arctic folds and wild-type Aβ42 filaments contain β-strands, like other amyloids. Thus, PiB-PET negativity of *APP*^E693G^ cases (28) does not reflect the absence of amyloid; it suggests instead that PiB does not recognise the fold of E22G Aβ. The same may be true of the Osaka mutation (33). It remains to be determined how these structures relate to what has been referred to as ‘protofibrils’ based on assembly experiments of synthetic E22G Aβ40 (24).

Tau filaments from case *AβPParc1* were identical to PHFs from sporadic and familial cases of AD (9,10). The same tau filament structures have been described in prion protein amyloidosis (13), as well as in familial British and Danish dementias (29). These findings are consistent with the suggestion that the Alzheimer fold of assembled tau is present whenever extracellular amyloid deposits form, irrespective of their structures and composition.

We showed previously that Aβ filaments from mice of the *App*^NL-F^ knock-in line, which express wild-type Aβ, are identical to those from human brains (35). Aβ filaments from the brains of mice of the *App*^NL-G-F^ knock-in line, which are made of two identical mutant protofilaments extending from D1-G37, differ from both wild-type and Arctic mutation filaments from human brains, raising doubts about the relevance of this mouse model for studying human disease.

The reasons for the differences in structure are unknown. It has been reported that murine BACE1 only cleaves human APP at position +1, whereas human BACE1 cleaves it at positions +1 and +11 (6). Another difference is that 100% of Aβ is mutant in knock-in mice (26), whereas the Arctic mutation is heterozygous and dominantly inherited (24). In the murine Arctic fold, the main chain conformation at G22 is incompatible with non-glycine residues, unlike in human Arctic folds, where other residues can be accommodated at this site with only minor conformational changes. This suggests that the incorporation of wild-type Aβ may inhibit formation and/or growth of mutant filaments with the murine Arctic fold, but not with the human Arctic folds.

Knock-in (26) and transgenic mouse models (8) have shown that the Arctic mutation is highly fibrillogenic when compared to wild-type Aβ. The fibrillation-promoting effects of E22G Aβ are the same for murine and human Arctic folds, namely increased hydrogen bonding between adjacent Aβ molecules and reduced electrostatic repulsion.

In summary, we report the structures of Aβ filaments from a case of AD with the Arctic mutation and from mouse knock-in line *App*^*NL-G-F*^. The murine Arctic fold is different from the human Arctic folds. Knowledge of the Aβ folds in human disease will inform the rational design of compounds that bind specifically to these filaments and the development of more relevant models for AD using *in vitro* assembly, cells and animals.

## ACKNOWLEDGEMENTS

This work was supported by the Electron Microscopy Facility of the MRC Laboratory of Molecular Biology. We thank Jake Grimmett, Toby Darling and Ivan Clayson for help with high-performance computing. We acknowledge Diamond Light Source for access and support of the cryo-EM facilities at the UK’s Electron Bio-imaging Centre (under proposal bi23268), funded by the Wellcome Trust, the MRC and the Biotechnology and Biological Sciences Research Council (BBSRC). This work was supported by the Medical Research Council, as part of the United Kingdom Research and Innovation (also known as UK Research and Innnovation) [MRC file reference number xxx]. For the purpose of open access, the MRC Laboratory of Molecular Biology has applied a CC BY public copyright licence to any Author Accepted Manuscript arising. M.G. is an Associate Member of the UK Dementia Research Institute.

## AUTHOR CONTRIBUTIONS

B.G., C.G., A.K. and A.N. identified the patient and performed genetic analysis and neuropathology; T.S. and T.C.S. provided *App*^NL-G-F^ knock-in mice; J.M. and I.L. organized breeding and characterized mouse tissues; Y.Y., M.H., S.L. and S.Y. P.-C. prepared filaments and performed mass spectrometry; Y.Y. and M.S. performed cryo-EM data acquisition; Y.Y., W.Z., A.G.M. and S.H.W.S. performed cryo-EM structure determination; M.G. and S.H.W.S. supervised the project and all authors contributed to the writing of the manuscript.

## FUNDING

This work was supported by the UK Medical Research Council (MC_UP_A025-1013 to S.H.W.S. and MC_U105184291 to M.G.) and the

US National Institutes of Health (UO1-NS110457).

## DECLARATIONS

### Ethics approval and consent

Studies carried out at Indiana University and the Karolinska Institutet (2011/962-31/1) were approved through the ethical review processes at each Institution’s Review Board. Informed consent was obtained from the patient’s next of kin. This study was approved by the Cambridgeshire 2 Research Ethics Committee (09/H0308/163).

### Data and materials availability

Cryo-EM maps have been deposited in the Electron Microscopy Data Bank (EMDB) with the accession numbers EMDB 16022, 16023 and 16027. Corresponding refined atomic models have been deposited in the Protein Data Bank (PDB) under accession numbers 8BFZ, 8BG0 and 8BG9. Please address requests for materials to the corresponding authors.

## SUPPLEMENTARY FIGURE LEGENDS

**Figure S1.**
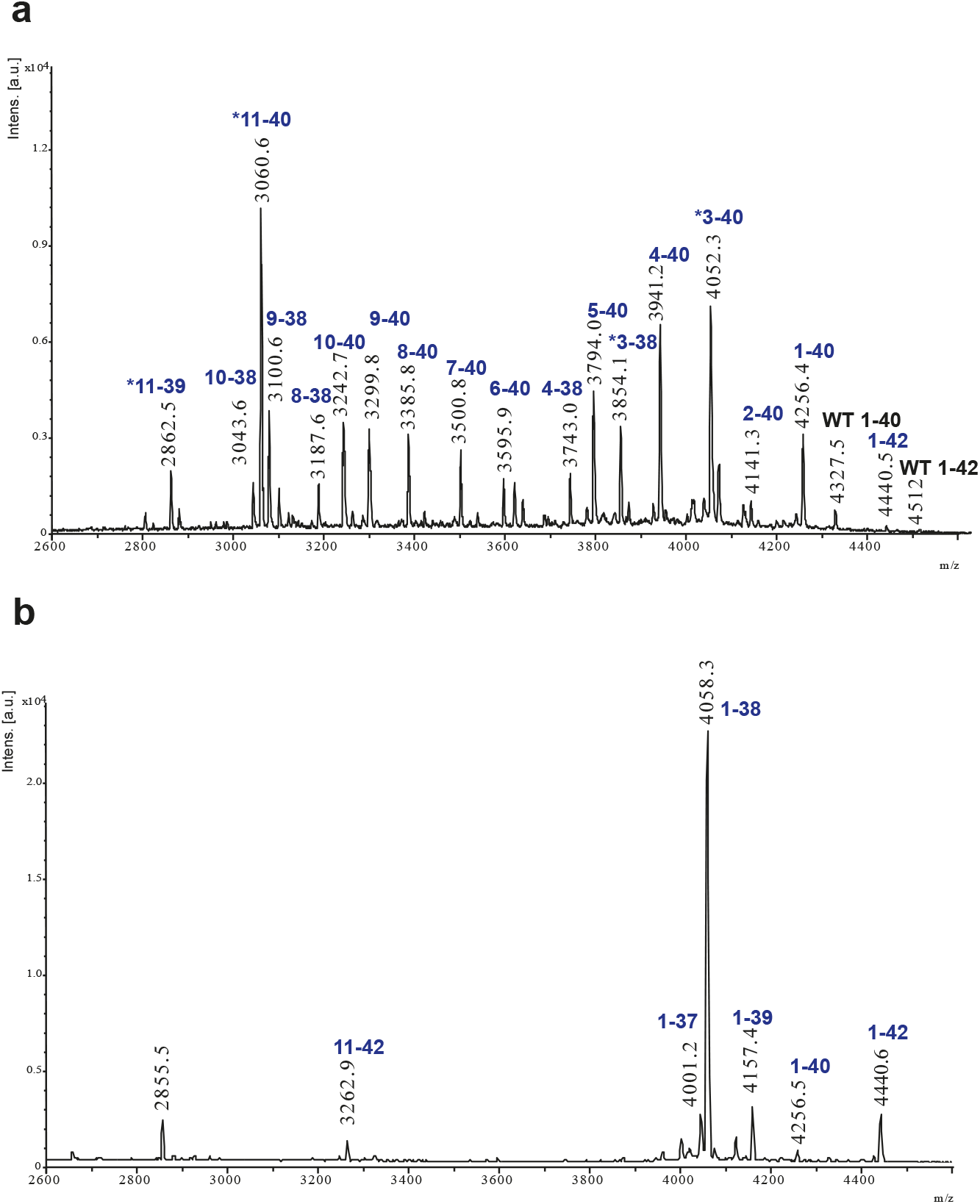
Mass spectrometric analysis. a. MALDI mass spectra for Aβ from the sarkosyl-insoluble fractions used for cryo-EM of human E22G Aβ. Mutant Aβ (blue), wild-type Aβ (black). Starred blue peptides are pyroglutamate-modified. b. MALDI mass spectra for Aβ from the sarkosyl-insoluble fractions used for cryo-EM of *App*^NL-G-F^ mouse brains. Mutant Aβ in blue.

**Figure S2.**
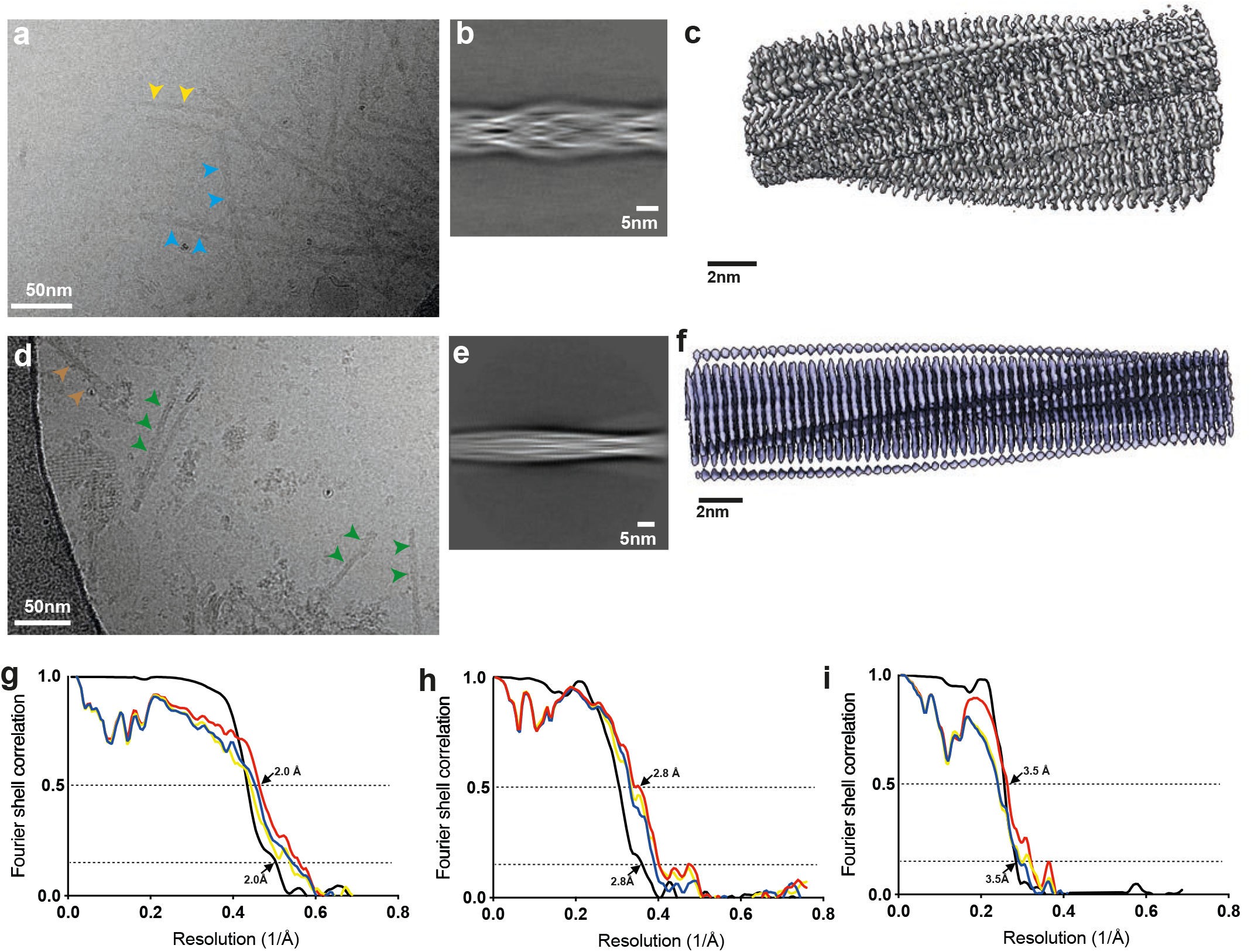
Cryo-EM micrographs and processing details. a,d. Cryo-EM micrographs of E22G Aβ filaments from temporal cortex of case *AβPParc1* (a) and E22G Aβ filaments from *App*^NL-G-F^ mouse brains (d). In (a), human Arctic tetrameric filaments are indicated with cyan arrows, dimeric type II Aβ42 filaments are indicated with orange arrows. In (d), dimeric filaments with the murine Arctic fold are indicated with green arrows and tetrameric filaments with the murine Arctic fold are indicated with brown arrows. Scale bar, 50 nm. b,e. 2D class averages of E22G Aβ filaments from human brain (b) and E22G Aβ filaments from *App*^NL-G-F^ mouse brains (e). c,f. 3D reconstructions of E22G Aβ filaments from human brain (c) and E22G Aβ filaments from *App*^NL-G-F^ mouse brains (f). Scale bar, 2 nm. g,h,i. Fourier shell correlation (FSC) curves for cryo-EM maps and structures of human Arctic tetrameric filaments (g), human Arctic dimeric filaments (h) and mouse *App*^NL-G-F^ Arctic dimeric filaments (i). FSC curves for two independently refined cryo-EM half maps are shown in black; for the final refined atomic model against the final cryo-EM map in red; for the atomic model refined in the first half map against that half map in blue; and for the refined atomic model in the first half map against the other half map in yellow.

**Table S1.**
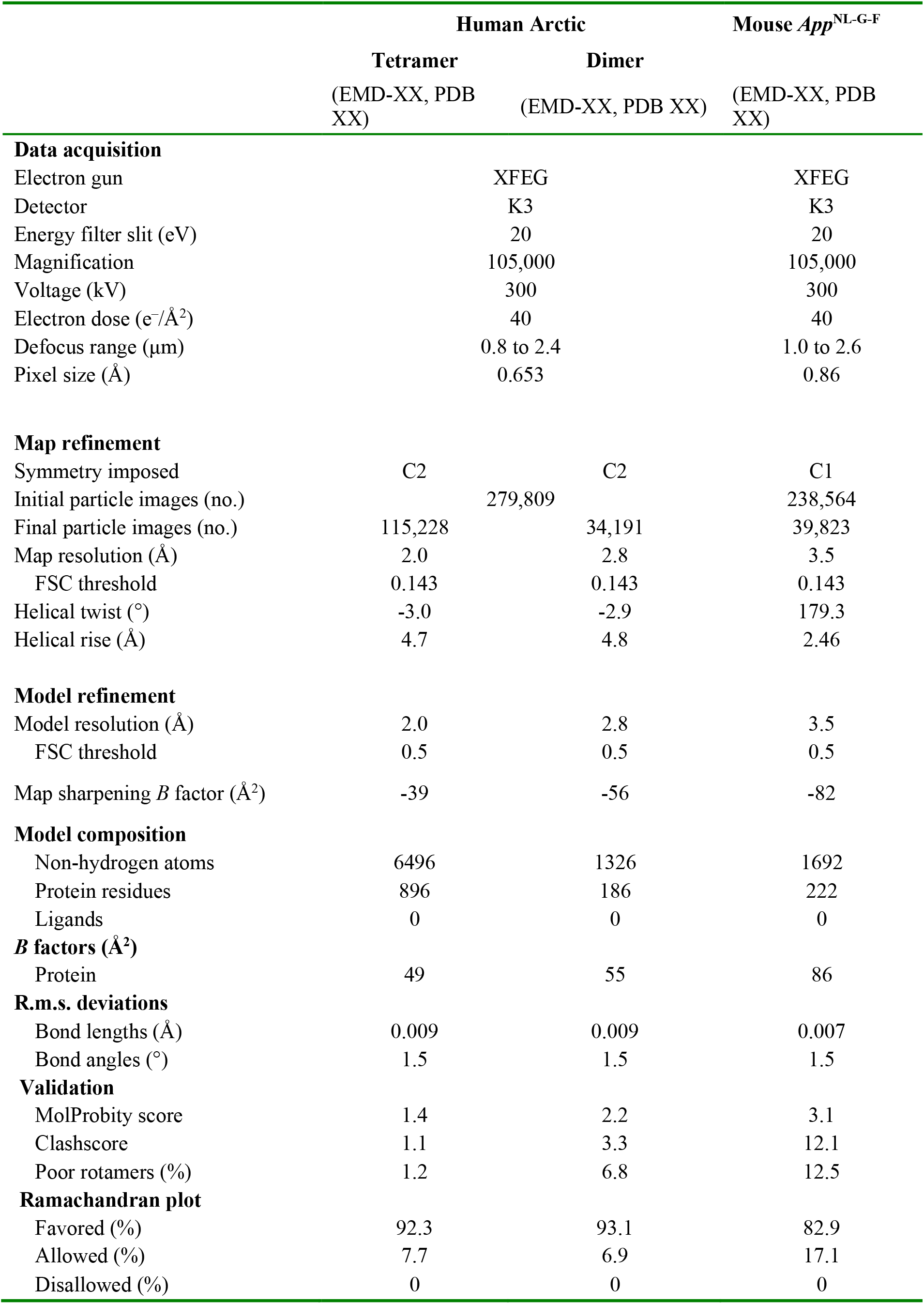
Cryo-EM data acquisition and structure determination.

